# Ebola virus requires phosphatidylserine scrambling activity for efficient budding and optimal infectivity

**DOI:** 10.1101/2020.03.16.994012

**Authors:** Marissa D. Acciani, Maria F. Lay-Mendoza, Katherine E. Havranek, Avery M. Duncan, Hersha Iyer, Olivia L. Linn, Melinda A. Brindley

## Abstract

**Abstract:** Ebola virus (EBOV) interacts with cells using two categories of cell surface receptors, C-type lectins and phosphatidylserine (PS) receptors. PS receptors typically bind to apoptotic cell membrane PS and orchestrate the uptake and clearance of apoptotic bodies. Many viruses coated with PS-containing lipid envelopes, acquired during budding from host cells, can also exploit these receptors for internalization. PS is restricted to the inner leaflet of the plasma membrane in homeostatic cells, an orientation that would be unfavorable for PS receptor-mediated uptake if conserved on the viral envelope. Therefore, it is theorized that viral infection induces host cell PS externalization to the outer leaflet during replication. Cells have several membrane scramblase enzymes that enrich outer leaflet PS when activated. Here, we investigate two scramblases, TMEM16F and XKR8, as possible mediators of cellular and viral envelope surface PS levels during recombinant VSV/EBOV-GP replication and EBOV virus-like particle (VLP) production. We found that rVSV/EBOV-GP and EBOV VLPs produced in XKR8 knockout cells contain decreased levels of PS in their outer leaflets. ΔXKR8-made rVSV/EBOV-GP is 70% less efficient at infecting cells through apoptotic mimicry compared to viruses made in parental cells. Our data suggest that virion surface PS acquisition requires XKR8 activity, whereas TMEM16F activity is not essential. Unexpectedly, we observed defective rVSV/G, rVSV/EBOV-GP, and EBOV VLP budding in ΔXKR8 cells, suggesting that phospholipid scrambling via XKR8 enhances both Ebola infectivity and budding efficiency. Overexpression of XKR8 dramatically increased budding activity, suggesting outer leaflet PS is required for both particle production and increased infectivity.

**Importance:** The Democratic Republic of the Congo experienced its deadliest Ebola outbreak from 2018 to 2020, with 3,444 confirmed cases and 2,264 deaths (as of March 12, 2020). Owing to the extensive damage that these outbreaks have caused in Africa, as well as its future epidemic potential, Ebola virus (EBOV) ranks among the top eight priority pathogens outlined by the WHO in 2018. A comprehensive understanding of Ebola entry pathways into target cells is critical for antiviral development and outbreak control. Thus far, host-cell scramblases TMEM16F and XKR8 have each been named as the sole mediator of Ebola envelope surface phosphatidylserine (PS). We assessed the contributions of these proteins using CRISPR knockout cells and two EBOV models: rVSV/EBOV-GP and EBOV VLPs. We observed that XKR8 is required for optimal EBOV envelope PS levels, PS receptor engagement, and particle budding across all viral models, whereas TMEM16F did not play a major role.

## Introduction

Ebola virus (EBOV) is an enveloped, non-segmented, negative-sense RNA filovirus that can cause severe hemorrhagic fever disease in those infected. Africa has endured multiple devastating EBOV outbreaks since the virus’s identification in 1976, of which the most severe, occurring in 2014-2015 in West Africa, led to more than 28,000 cases and 11,000 deaths (1). Most recently, the Democratic Republic of the Congo experienced an EBOV outbreak spanning from August of 2018 to February 2020, the country’s 10th outbreak in the past 40 years, which was declared a public health emergency of international concern by the World Health Organization (2). Although the rVSV-ZEBOV-GP vaccine was deployed and licensed in December of 2019, the DRC outbreak took more than a year to contain (3–6). With case fatality rates historically reaching up to 90%, continuous virus reemergence, and mounting evidence of viral persistence in EBOV survivors, the risk of further high-mortality-associated EBOV outbreaks remains (7, 8).

EBOV transmission occurs via direct contact of the virus with broken skin or mucous membranes, where virions encounter their initial target cell types (myeloid dendritic cells and macrophages) (9, 10). EBOV enters these cells through a dynamin-dependent macropinocytosis pathway (11, 12), followed by cathepsin-EBOV glycoprotein (GP) proteolysis, GP interaction with endosomal receptor Niemann–Pick C1 (NPC1), and virus-cell membrane fusion within the endosome (13, 14). EBOV’s broad cell tropism has been attributed, in part, to its use of a variety of cell-surface receptors. C-type lectins, T-cell immunoglobulin and mucin domain (TIM) proteins, and Tyro-3/Axl/Mer (TAM) receptor tyrosine kinases can facilitate EBOV cell-surface attachment and internalization (15–21). TIM and TAM family proteins share a common ligand, an anionic phospholipid with a serine head group called phosphatidylserine (PS). Professional and non-professional phagocytes expressing TIMs or TAMs use these receptors to detect and clear apoptotic bodies, which are abnormally rich in plasma membrane outer leaflet PS. Viral envelopes also contain outer leaflet PS and can, therefore, gain access to clearance cells by binding to their PS receptors, a viral entry strategy referred to as “apoptotic mimicry” (22). Several studies have confirmed the importance of PS receptor-mediated EBOV entry in cultured cells (17, 18, 20–23). Futhermore, groups examining EBOV pathogenesis in TIM^−^/^−^ mice found TIM1 to be critical for viral loads in the liver, kidneys, spleen, and lymph nodes, and serum cytokine induction, which are both characteristic of human EBOV infection (24, 25).

Many pathogenic enveloped viruses can infect cells through apoptotic mimicry, including EBOV (17, 18, 20–23), Lassa virus (26–28), dengue virus (23, 29–31), Zika virus (32), and vaccinia virus (33). These viruses obtain PS from host cells during viral budding, wherein virions are enshrouded in a portion of lipid bilayer from an internal organelle membrane or the plasma membrane (34–37). EBOV buds through the host-cell plasma membrane, thereby acquiring an envelope with a lipid composition reflecting that of the cellular surface (34). In healthy cells, plasma membrane phospholipid distribution is highly asymmetric, with the majority of PS restricted to the inner leaflet (38). Because this orientation would be unfavorable for PS receptor engagement if transferred to the viral envelope, it has been theorized that viral infection induces host-cell phospholipid scrambling, resulting in viral progeny with envelopes rich in externalized PS.

Cell PS externalization is mediated by transmembrane scramblases that nonspecifically transfer phospholipids between plasma membrane bilayer leaflets, reversing lipid asymmetry (39). Of all the identified proteins demonstrating phospholipid scramblase activity, TMEM16F and XKR8 are the most prominent and well characterized (40). TMEM16F activity is reversible and is stimulated by large increases in cellular cytosolic calcium (41–44), whereas XKR8 is permanently activated during apoptosis, specifically by caspase-3/7 cleavage (45–47). In recent publications by Nanbo et al. (48) and Younan et al. (49), the authors proposed that XKR8 and TMEM16F are required for Ebola envelope surface PS acquisition. Younan et al. observed widespread calcium-induced TMEM16F-dependent cellular PS scrambling in Huh7 cells following wild-type (WT) EBOV infection, and that knocking down TMEM16F reduced EBOV surface PS and virus cell–cell spread. Knocking down XKR8 did not inhibit EBOV-induced PS scrambling. Alternatively, Nanbo et al. attributed Ebola virus-like particle (VLP) envelope PS levels and subsequent VLP internalization efficiency to XKR8 activation. This group further suggested that the expression of EBOV VP40 and GP proteins at the plasma membrane of Vero E6 cells induces localized caspase cleavage and XKR8-mediated PS scrambling at EBOV VLP budding sites.

In this study, we sought to clarify the contributions of TMEM16F and XKR8 to EBOV envelope surface PS levels by producing either recombinant vesicular stomatitis virus containing the Zaire EBOV glycoprotein (rVSV/EBOV-GP) or EBOV virus-like particles (VLPs) in human haploid (HAP1) cells knocked-out for TMEM16F (ΔTMEM16F cells) and XKR8(ΔXKR8 cells). We compared rVSV expressing its native glycoprotein (rVSV/G) or rVSV/EBOV-GP in parental, ΔTMEM16F, and ΔXKR8 cells. rVSV/EBOV-GP cell–cell spread, virion specific infectivity, and viral envelope PS levels were reduced when the virus was produced in ΔXKR8 cells. EBOV VLP surface PS was similarly dependent on XKR8 activity. Additionally, we observed reduced rVSV budding efficiencies in ΔXKR8 cells that repeated in the VLP model. This suggest that PS scrambling activity is needed not only for viral surface PS, but also for viral particle assembly and/or egress.

## Materials and Methods

### Cell lines and transfections

HAP1, HAP1ΔXKR8 (HZGHC005916c007), and HAP1ΔTMEM16F (HZGHC002956c003) cells (Horizon Discovery) were maintained in Iscove’s media supplemented with 10% (vol/vol) fetal bovine serum (FBS). Knock-out HAP1 cells were verified by amplifying the exon 2 of XKR8 and confirming it contained a two base pair insertion causing a frameshift mutation. ΔTMEM16f exon 5 lacked 20 base pairs. Vero (African green monkey kidney) cells stably expressing human SLAM and baby hamster kidney (BHK-21) cells stably expressing T7 polymerase (BSR-T7/5 [BHK-T7]) (50) were maintained in Dulbecco’s modified Eagle’s medium (DMEM) supplemented with 5% (vol/vol) FBS (51). All cells were kept at 37°C and 5% CO_2_. BHK-T7 transfections were performed using Gene Juice (MilliporeSigma) and HAP1 and Vero transfections used JetOptimus (Polyplus) as per manufacturer’s instructions.

### Drugs and compounds

For apoptosis induction, HAP1 cells were treated with 1 μM MG132 (Enzo Life Sciences) in Iscove’s medium for 24 hours at 37°C and 5% CO_2_ prior to annexin V staining. To induce the release of intracellular calcium stores, cells were treated for 5 minutes with 5 μM A23187 (Enzo Life Sciences) in PBS at room temperature prior to staining. To functionally inhibit XKR8, VLP-producing cells were treated with 20 μM Z-VAD-FMK (Enzo Life Sciences) 3 hours following VLP transfection.

### Annexin V cell staining

Phosphatidylserine externalization in HAP1-derived cells was quantified by co-staining with both annexin V-Pacific Blue (AnV-PacBlue) (Invitrogen) and propidium iodide (PI) (Sigma Aldrich). Cells were lifted using trypsin without EDTA, washed with PBS, and incubated in 100 μL of annexin binding buffer (10 mM HEPES, 140 mM NaCl, and 2.5 mM CaCl2, pH 7.4) containing AnV-PacBlue and PI for 30 minutes on ice. Cells were then diluted 1:4 in annexin binding buffer and analyzed in a flow cytometer (BD-LSRII). Cell populations were gated using forward scatter/side scatter. We excluded doublets and cell aggregates by gating cell populations using side scatter area/side scatter height. The same gates were used for all samples. Mean Fluorescence Intensity (MFI) of the Pacific Blue fluorophore in PI negative cells was quantified and averaged over a minimum of 3 independent experiments.

### Immunoblot analysis

Parental and scramblase knockout HAP1 cells were subjected to SDS-PAGE and immunoblot analysis using anti-TMEM16F (1:1000, Sigma Aldrich, HPA038958) and β-actin C4 (1:1000, Santa Cruz Biotechnology Inc., sc-47778) antibodies. Immunoblots were probed with appropriate secondary antibodies conjugated with HRP and imaged with a ChemiDoc XRS digital imaging system (Bio-Rad).

### Viruses

Measles virus (MeV-GFP), Edmonston strain, expressing GFP in a pre-N position has been previously described (52).

### Generation of replication competent recombinant VSV

The full-length cDNA clone of the VSV genome (Indiana serotype) in which a multiple cloning site (MCS) (5’-MluI-XmaI-SmaI-EagI-NheI-3’) replaced the glycoprotein and GFP is encoded as an additional transcriptional unit between the MCS and the L gene, pVSV-ΔG-GFP-2.6 (courtesy of Dr. Michael Whitt; KeraFAST #EH1027) (53), was used as the backbone for all recombinant viruses. The VSV G Indiana strain (accession number: NP_041715.1) and EBOV glycoprotein Zaire strain (accession number: AAA96744.1) sequences were PCR amplified (ATG-stop codon) with primers that added a 5’ MluI and 3’ NheI sites (54). The glycoproteins were cloned into the genome using the MluI and NheI restriction sites. Viral constructs containing nano-luciferase within the viral particle were made by replacing the matrix (M) gene with the M-nLuc construct. M-nLuc was generated by adding the nano-luciferase coding region after residue 37 as was previously done with GFP (55).

Viruses were recovered by co-transfecting BHK-T7 cells with a plasmid carrying a cDNA copy of the recombinant VSV genome and plasmids encoding VSV polymerase (L), nucleocapsid (N), phosphoprotein (P), and glycoprotein (G) proteins (courtesy of Dr. Michael Whitt; KeraFAST) (53). All constructs were under the control of the T7 promoter. Transfected BHK-T7 cells were overlaid on Vero cells after 48 hours, and virus stocks were passaged onto fresh Vero cells using a low MOI 0.001 to generate passage 2 and 3 stocks. All experiments were completed with virus produced from passage 2 or 3 in Vero cells. Viral titers were determined by performing end-point dilutions on Vero or HAP1 cells and calculating the 50% tissue culture infective dose (TCID_50_) according to the Spearman-Karber method (52).

### Multi-step growth curve

HAP1, ΔXKR8, and ΔTMEM16F cells were seeded in 12-well plates (1.25×10^5^ cells/mL for rVSV/EBOV-GP or 2.5×10^5^ cells/mL for rVSV/G and MeV) and infected with the indicated virus at an MOI of 0.01. Virus-containing media was removed 1 hour following infection and replaced with fresh Iscove’s medium. At each time point, the supernatant was collected and replaced with fresh Iscove’s medium. Because MeV is cell associated, the infected monolayer was scraped in 500 μL of supernatant and viral particles were released with two freeze-thaw cycles (52). All samples were stored at −80°C and titrated on Vero cells.

### qRT-PCR

Viral RNA was isolated from infected cell supernatant (QIAamp® Viral RNA Mini Kit, Qiagen) and cell lysate (RNeasy Mini Kit, Qiagen). All samples were collected in duplicate. RNA was reverse transcribed into cDNA (High-Capacity RNA-to-cDNA™ Kit, Thermo Scientific). VSV genome copies were measured with qRT-PCR using the TaqMan Gene Expression Master Mix (Applied Biosystems), VSV-L forward and reverse primers, and probe (TaqMan MGB Probe) (56). Each sample was analyzed in duplicate, and each assay contained a DNA standard curve (VSV molecular clone, ranged from 3 to 3×10^6^ plasmid copies), no template, and no primer controls. Because the primers detect VSV-L RNAs, copies generated from cellular RNA detected both full-length genome as well as VSV-L transcripts. We extrapolated VSV copy numbers from the generated standard curve using the Applied Biosystems protocol. Final copy numbers were adjusted by back-calculations to the total RNA and cDNA volume and expressed as copies per lysate or supernatant sample.

### Virus Entry and Replication: qRT-PCR

HAP1 and ΔXKR8 cells were seeded in 12-well plates (5×10^5^ cells/mL). After 24 hours, cells were infected with rVSV/G (MOI= 0.01) or rVSV/EBOV-GP (MOI= 0.05) for 2 hours at 37°C 5% CO_2_. Virus-containing media was removed, and cells were treated with citric acid buffer (40 mM citric acid, 10 mM KCl, 135 mM NaCl, pH 3.0) for 1 minute to inactivate any remaining viral particles. Citric acid was removed, cells were washed with Iscove’s media, and fresh Iscove’s media was added.

#### Viral entry

To examine virus entry, viral genomes were quantified in cells 2 hours after infection. Viral copy number was determined by comparing the C(t) values in the total cellular RNA to a DNA standard curve. Three independent trials were performed.

#### Viral RNA replication

Viral genome copy numbers were quantified in cells and supernatants 12 hours post-infection. Cell viral copy numbers were adjusted for total ng RNA isolated and represented viral replication ability. Samples were collected in duplicate from 3 independent trials.

### GFP Virus Entry and spread

#### GFP virus entry

HAP1 and ΔXKR8 cells were seeded in 48-well plates (5×10^5^ cells/mL). After 24 hours, cells were incubated with rVSV/G (MOI= 2) at 4°C for 45 minutes. Virus-containing media was removed, cells were washed with PBS, and fresh Iscove’s medium was added. We observed that rVSV/EBOV-GP bound and infected HAP1 cells approximately 100-fold less efficiently than rVSV/G, thus a HAP-specific titer and MOI were used for all future experiments. We also infected HAP1 cells with rVSV/EBOV-GP for 2 hours (MOI= 20) at 37°C, then removed the virus-containing media, and fresh Iscove’s medium was added. rVSV/G and rVSV/EBOV-GP-infected cells were trypsinized 16 and 12 hours after infection respectively, fixed with 4% formaldehyde, and GFP-positive virus-infected cells were quantified using flow cytometry (BD-LSRII).

#### GFP virus cell-cell spread

Cells were seeded in 12-well plates (5×10^5^ cells/mL). After 24 hours, cells were incubated with rVSV/G (MOI= 0.005) or rVSV/EBOV-GP (MOI= 0.01) for 2 hours at 37°C, inoculum was removed, and fresh media was replaced. Representative images were captured using the Zoe microscope (Bio-Rad) (magnification x20) at the indicated time points.

### rVSV budding assays

#### Luminescence assay

Cells were seeded in 12-well plates (5×10^5^ cells/mL). After 48 hours, cells were incubated with rVSV-MnLuc/G (MOI= 0.1) or rVSV-MnLuc/EBOV-GP (MOI= 0.25) for 2 hours at 37°C, inoculum was removed, cells were citric acid treated, washed, and fresh media was replaced. Twelve hours post infection the cell lysate and supernatants were collected, and assayed for nano-luciferase activity using Nano-Glo luciferase substrate (Promega) and luminescence was measured in the Glomax® Explorer (Promega). Budding efficiency was calculated by dividing the signals from the supernatant by the cell lysate. In all experiments, HAP1 budding efficiency was set to 100%.

#### Protein production

HAP1 and ΔXKR8 cells were seeded in 12-well plates (5×10^5^ cells/mL). After 24 hours, cells were infected with rVSV/G (MOI= 0.05) or rVSV/EBOV-GP (MOI= 4) for 2 hours at 37°C 5% CO_2_. Virus-containing media was removed, cells were treated with citric acid buffer for 1 minute, washed with Iscove’s media, and fresh Iscove’s media was added. Cells were incubated at 37°C 5% CO_2_ for 12 hours, after which cell lysates and supernatants were collected. Cell lysates and supernatants were subjected to immunoblot analysis for antibodies against cellular LAMP-1 (1:2000, SouthernBiotech, 9835-01), VSV-G (1:2000, Apha Diagnostic, VSIG11-S), EBOV-GP (1:1000, IBT Bioservices, 0201-022), and VSV-M (1:2500, Kerafast, EB0011).

### Transmission Electron Microscopy

Confluent flasks of HAP1 and ΔXKR8 cells were infected with rVSV/EBOV-GP (MOI= 1) and incubated for 2 days. Cells were fixed with 2.5% glutaraldehyde and 2% paraformaldehyde in PBS for 1 hour at room temperature. Cells were rinsed several times in PBS and enrobed in 4% Noble agar. Enrobed samples were rinsed several times in PBS and incubated for 1 hour in 1% OsO_4_ in PBS, and rinsed again several times in deionized water. Samples were then dehydrated in a series of graded ethanol. Subsequently, samples were infiltrated with Embed 812 resin and processed for 1-2 days at 60°C. Thin sections were mounted on copper mesh grids and post-stained with uranyl acetate and lead citrate. Specimens were viewed on a JEOL JEM1011 transmission electron microscope (JEOL, Inc., Peabody, MA). Samples were produced three independent times and representative images are shown.

### Viral Particle Specific Infectivity

HAP1 and ΔXKR8 cells were seeded in T75 flasks (5×10^5^ cells/mL). After 24 hours, cells were infected with rVSV/G (MOI= 0.01) or rVSV/EBOV-GP (MOI= 2) for 2 hours at 37°C and 5% CO_2_. Virus-containing media was removed, cells were treated with citric acid buffer for 1 minute and washed, and fresh Iscove’s media was added. After 24 hours, the supernatant was collected, concentrated and purified using Vivaspin 20 ultrafiltration units (300kDa MW cut-off, Sartorius), and stored at −80°C.

#### Specific Infectivity

Specific infectivity for virus produced in HAP1 and ΔXKR8 cells was calculated by taking the ratio of genome copy numbers to PFU/mL for each sample. To quantify genome copy numbers in each sample, qRT-PCR was performed on harvested virus as was described above. The PFU/mL for all samples was determined using plaque assays on Vero cells. Specific infectivity values were averaged from virus produced from 3-4 independent trials.

#### Vero Specific Infectivity Rescue

Vero cells were seeded in a 24-well plate (5×10^5^ cells/mL) and infected with HAP1-made or ΔXKR8-made rVSV/G (MOI= 0.01) or rVSV/EBOV-GP (MOI= 0.05). Infection proceeded for 2 hours at 37°C and 5% CO_2_. Virus-containing media was removed, and cells were treated with citric acid buffer for 1 minute, washed, and fresh Iscove’s media was added. After 24 hours, the supernatants were collected and stored at −80°C. Specific infectivity for these samples was determined as described above.

### Viral Particle Glycoprotein Incorporation

HAP1 and ΔXKR8 cells were seeded in 6-well plates (5×10^5^ cells/mL). After 24 hours, cells were infected with rVSV/EBOV-GP (MOI 18) for 2 hours at 37°C 5% CO_2_. Virus-containing media was removed, cells were treated with citric acid buffer for 1 minute, washed, and fresh Iscove’s media was added. Cells were incubated at 37°C 5% CO_2_ for 24 hours, after which cell supernatants were collected. A portion of supernatants was used for viral genome quantification in order to normalize samples. Normalized supernatants were precipitated using 10% (wt/vol) TCA (57). The TCA-treated proteins were pelleted (17,000 × *g*, 30 min, 4°C), washed with acetone, dried, and denatured using SDS-urea buffer (200 mM Tris [pH 6.8], 8 M urea, 5% SDS, 0.1 mM EDTA, 0.03% bromophenol blue). The denatured TCA-treated proteins were subjected to immunoblot analysis for VSV M and EBOV GP protein levels using anti-VSV M (1:2500, Kerafast, EB0011) and anti-EBOV GP (1:1000, IBT Bioservices, 0201-022) antibodies.

### rVSV Surface PS Quantification

HAP- or ΔXKR8-produced purified rVSV/G or rVSV/EBOV-GP (1.5×10^9^ genomes) were conjugated to 4-μm aldehyde/sulfate latex beads (Thermo Fisher Scientific) overnight at 4°C with gentle shaking. Beads were blocked with 1% BSA in PBS for 2 hours while rotating at room temperature. The reaction was quenched by adding 100 mM of glycine for 30 minutes at room temperature. Beads were washed 3 times with 1% BSA in PBS, followed by a 15 minute incubation in 100 μL of AnV binding buffer containing AnV-PacBlue at room temperature. Beads were diluted 1:5 in AnV binding buffer and analyzed using flow cytometry. Beads were then washed 3 times with 1% BSA in PBS, followed by a 2 hour incubation with antibodies against VSV-G (1:200, Alpha Diagnostic International, VSIG11-S) or EBOV-GP (1:50, IBT Bioservices, 21D10) at room temperature with gentle shaking. Beads were then washed and incubated with anti-rabbit FITC (1:100, KPL, 02-15-16) or anti-mouse APC (1:50, Jackson ImmunoResearch, 115-135-164) secondary antibodies for 1 hour at room temperature with gentle shaking. Samples were analyzed by flow cytometry (BD-LSRII).

### Generation of EBOV VLPs

The Zaire EBOV GP gene was expressed using a pcDNA vector under control of the CMV promoter (Invitrogen, Carlsbad, CA) (54, 58). The pCAGGS-NP-EBOV plasmid encoding the Zaire EBOV NP gene was a gift from Elke Mühlberger (Addgene plasmid #103049) (59). EBOV mCherry-VP40 was a gift from Judith White (Addgene plasmid #74421) (60). We replaced the mCherry reading frame with nano-luciferase (nLuc-VLP) (Promega) or codon optimized GFP (GFP-VLP). The XKR8 reading frame was cloned into pcDNA-intron (61) and a 3xFLAG tag was added to the N-terminus. VLPs were produced by co-transfecting EBOV VP40, NP, and GP at a 1:1:1 ratio. In some experiments additional XKR8_FLAG_ or empty vector was also transfected.

#### VLP budding

HAP1, ΔXKR8, or Vero cells were seeded in 24-well plates (5×10^5^ cells/mL). After 24 hours, cells were transfected with VP40-nLuc, NP, and GP plasmids, media was replaced 4 hours post transfection, and supernatants and cell lysates were collected 24 hours post transfection and cleared of cell debris with centrifugation. Luciferase activity was detected using Nano-Glo Luciferase substrate (Promega) and luminescence was measured in the Glomax® Explorer (Promega). In experiments that included DMSO or Z-VAD-FMK, treatments were added during media replacement 4 hours following transfection. Budding efficiency was determined by dividing the luminescence signals in the supernatant by the cell lysate. In all experiments, HAP1 budding efficiency was set to 100%.

#### EBOV VLP Surface PS Quantification

HAP1 and ΔXKR8 cells were seeded in 6-well plates (5×10^5^ cells/mL). After 24 hours, cells were transfected with VP40-GFP, NP, and GP, with or without XKR8_FLAG,_ or EBOV VP40-GFP only (to produce VLPs lacking GP). Media was replaced 4 hours after transfection. VLPs were harvested 24 and 48 hours after transfection and pelleted (17,000 x g for 1 hour at 4°C). Supernatant was removed, and pellets containing VLPs were resuspended in 100 μL PBS and stored at −80°C. For GP staining, VLPs were incubated with chimeric anti-EBOV GP mAb (h13F6) (1:500, IBT Bioservices 0201-022) for 1 hour, followed by anti-human Alexa Fluor647 (1:1000, Jackson ImmunoResearch, 109-605-098) for 30 minutes. VLPs were then diluted 1:5 in PBS. For AnV staining, VLPs were incubated with AnnexinV-PE conjugate (AnV-PE) (Invitrogen) in AnV binding buffer for 30 minutes at 4°C. VLPs were then diluted 1:5 in AnV binding buffer and analyzed by flow cytometry. VLP samples were analyzed using the small particle detector on the Acea Novocyte Quanteon 4025, gating for GFP positive particles.

### Statistical Analysis

Results are presented as averages with standard errors of the mean (SEM) from at least 3 independent trials. Averages were compared using the Student’s t-test or Welch’s t-test if unequal variance in JMP® Pro software (version 13.2.0) or GraphPad Prism. A p-value of less than 0.05 was considered to be statistically significant.

## Results

### XKR8 is required for apoptosis-induced PS externalization in HAP1 cells

HAP1 cells have been used to screen and identify cellular genes critical for EBOV entry, as these nearly haploid cells are straightforward to knock out genes and are susceptible to EBOV infection (13, 62). In order to investigate the specific contributions of TMEM16F and XKR8 scramblases to EBOV replication, ΔTMEM16F and ΔXKR8 knockout HAP1 cell lines were first tested for scramblase activity. XKR8 is activated during apoptosis by caspase cleavage; therefore, we treated parental and knockout cells with MG132, a membrane-permeable proteasome inhibitor that induces caspase-3 cleavage and apoptosis (63), and stained for PS in the outer leaflet using AnnexinV-PacificBlue (AnV-PacBlue) (Fig. 1A and B). MG132-treated HAP1 and ΔTMEM16F cells showed a marked increase in surface PS, whereas ΔXKR8 surface PS levels were significantly reduced, indicating that XKR8 is the primary mediator of apoptosis-induced PS scrambling in HAP1 cells. HAP1 cell lines were also treated with calcium ionophore A23187 to test for calcium-dependent scramblase activity (Fig. 1A and B). All three cell lines resulted in robust AnV staining, indicating that both scramblase deficient cells retain calcium-mediated PS scrambling activity. We performed immunoblot analysis on cell lysates and confirmed that no TMEM16F protein was detected in ΔTMEM16F cells (Fig. 1D). Thus, while XKR8 is a primary mediator of apoptosis-induced PS externalization in HAP1 cells, TMEM16F is not essential for calcium-induced PS externalization.

**Figure 1.**
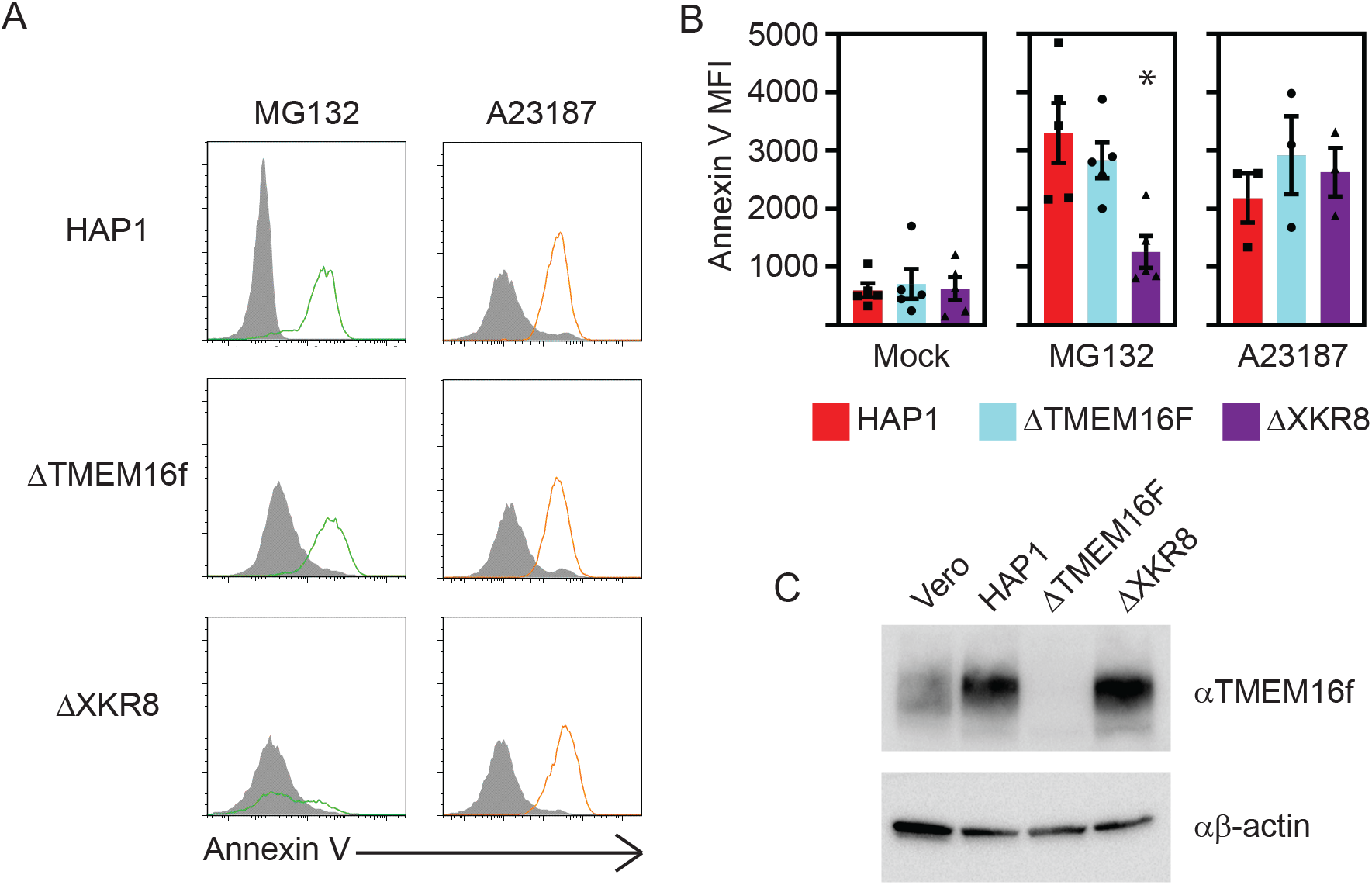
PS scrambling abilities of WT and KO HAP1 cells. HAP1, ΔTMEM16F, and ΔXKR8 cells were treated with apoptosis inducer MG132 (1μM) or calcium ionophore A23187 (5μM), double stained with AnV-PacBlue and PI, and analyzed using a BD LSRII flow cytometer. Anv-PacBlue fluorescence of live, single, non-necrotic cells are shown as (A) histograms and (B) average Mean Fluorescence Intensities (MFIs). C) Immunoblot probing for beta-actin and TMEM16F confirming the absence of TMEM16F in ΔTMEM16F cells. Values shown are averages of at least three independent experiments ± SEM, *p<0.05, student’s t-test.

### rVSV/EBOV-GP replication is impaired in ΔXKR8 cells

In order to elucidate the importance of scramblases in EBOV-GP-mediated replication, we performed multistep growth curves using rVSV encoding EBOV-GP (rVSV/EBOV-GP) or its native glycoprotein (rVSV/G), as well as measles virus (MeV) in parental and knockout HAP1 cell lines (Fig. 2). The internalization pathways of both rVSV/G and MeV are initiated by specific cell surface protein interactions and are not strongly associated with apoptotic mimicry (17, 64–66). All viruses were titrated on Vero-hSLAM cells that naturally produce PS receptors (TIM-1 and Axl) as well as MeV and rVSV-G receptors, but do not produce EBOV-enhancing C-type lectins (51, 67). Titers of rVSV/EBOV-GP produced in ΔXKR8 cells were significantly reduced compared to titers generated in HAP1 and ΔTMEM16f cells at all time points following infection (Fig 2A). The greatest differences occurred 72 hours post-infection, where we observed a 4.6-log reduction of infectious virions present in ΔXKR8 cell supernatant. However, rVSV/EBOV-GP titers in ΔXKR8 cells eventually recovered, suggesting that EBOV-GP-mediated replication is significantly delayed rather than strictly inhibited in these cells. rVSV/G spread similarly in all cell types, and although the titers in ΔXKR8 cells were consistently lower than those in HAP1 cells, they did not significantly differ at any time point (Fig. 2B). MeV also propagated through all cell lines with similar efficiencies (Fig. 2C), suggesting that cells lacking XKR8 scrambling activity are uniquely defective in entry or spread of EBOV-GP. Because we did not detect a functional defect in ΔTMEM16F cells with AnV staining, it was unsurprising that all viruses were able to propagate in these cells as efficiently as parental HAP1 cells.

**Figure 2.**
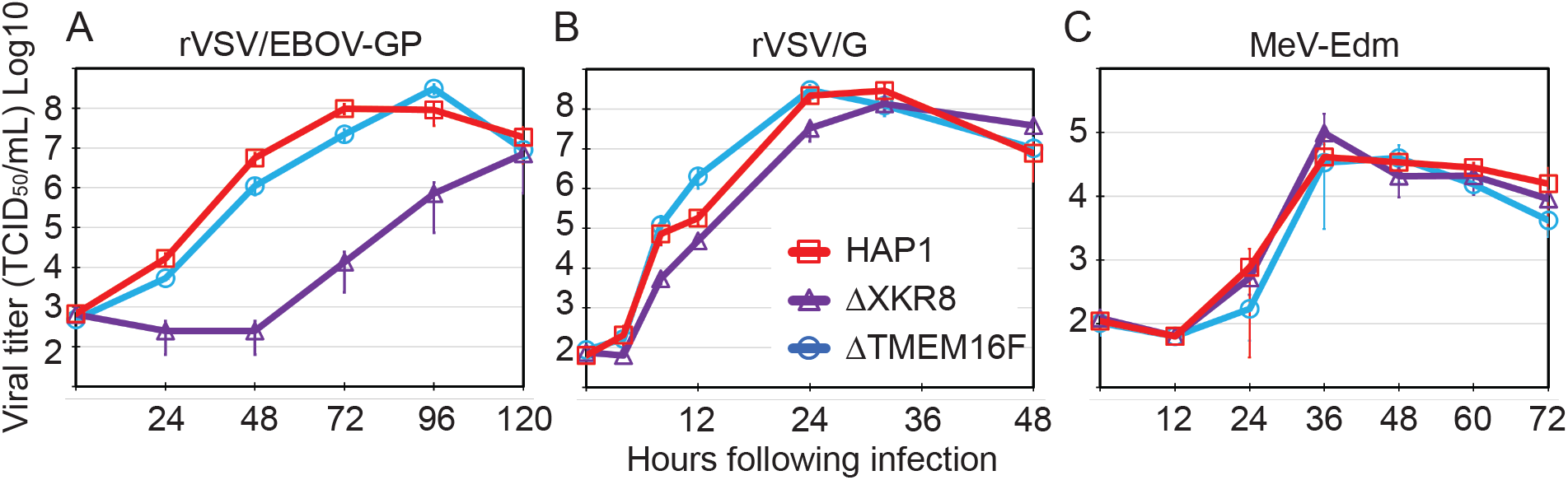
rVSV/EBOV-GP replication delayed in cells lacking XKR8. HAP1, ΔTMEM16F, and ΔXKR8 cells were infected with rVSV/EBOV-GP (A), rVSV/G (B), or MeV (C) (MOI= 0.01). Cell supernatants (A and B) or cell-associated virus (C) were collected at the indicated timepoints and titrated on Vero cells by performing end-point dilutions. Values shown are the average log10(TCID_50_/mL) of at least three independent experiments ± SEM.

### Knocking out XKR8 does not inhibit viral entry or genome replication

The rVSV/EBOV-GP replication delay observed in ΔXKR8 cells could be caused by inefficiencies at a number of steps in the virus replication cycle. Because the phenotype correlated with EBOV-GP, we examined the ability of ΔXKR8 cells to support early rVSV/EBOV-GP entry and one round of rVSV/EBOV-GP replication. XKR8 is inactive in homeostatic cells, therefore we did not expect XKR8 deletion to impact either of these stages in the viral life cycle. We quantified rVSV/EBOV-GP viral genomes internalized within HAP1 and ΔXKR8 cells following a 2◻hour entry period (Fig. 3A). We also infected cells with rVSV/EBOV-GP and measured the proportion of GFP-positive cells via flow cytometry after one round of replication (Fig. 3B). We were unable to detect significant entry differences between HAP1 and ΔXKR8 cells in either assay, indicating that the rVSV/EBOV-GP replication delay did not occur during initial infection.

**Figure 3.**
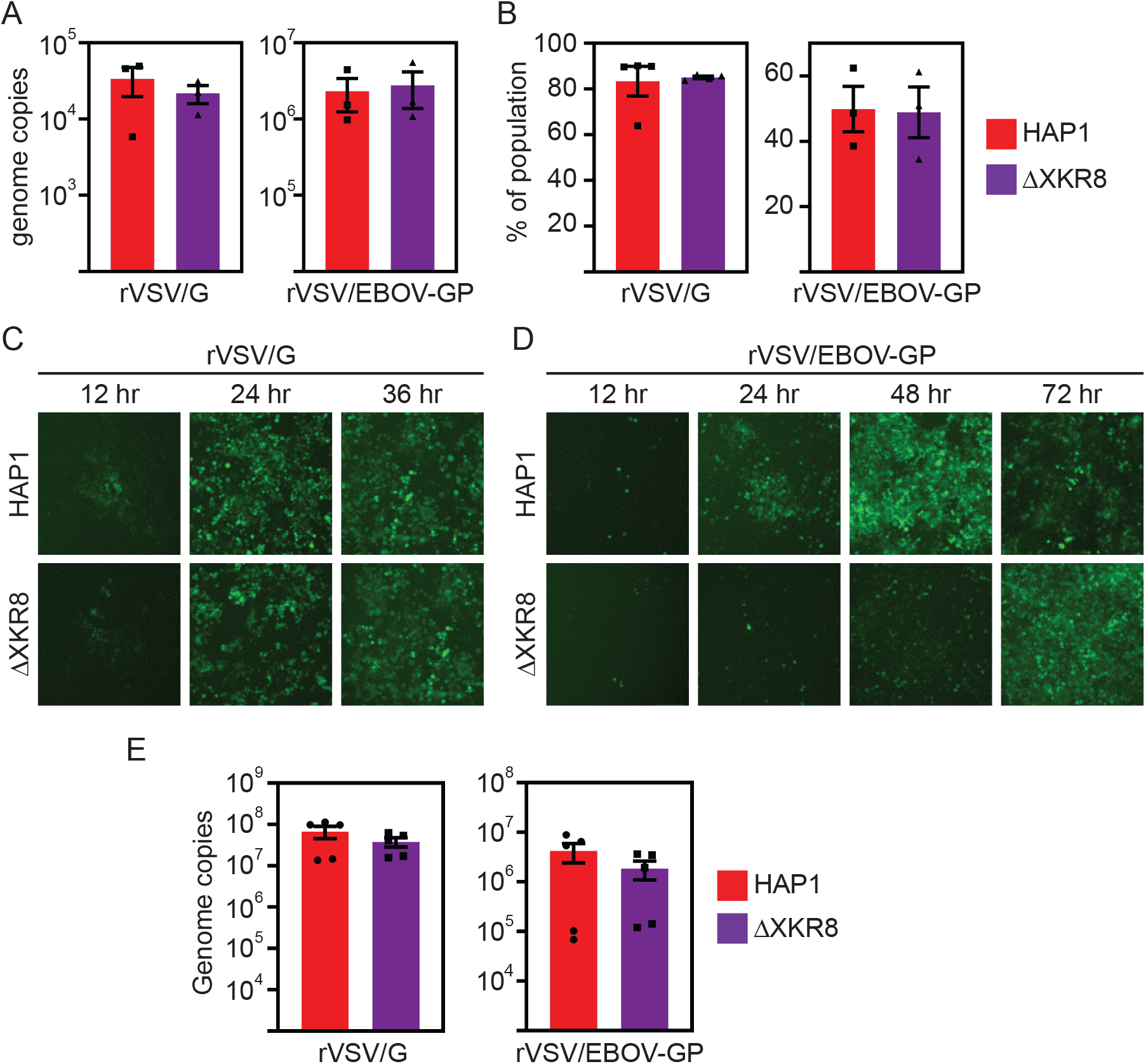
rVSV/EBOV-GP requires XKR8 for efficient cell-cell spread, but it is not required for entry or replication. A) HAP1 and ΔXKR8 cells were infected with rVSV/G (MOI= 0.01) or rVSV/EBOV-GP (MOI 0.05). After 2 hours, early viral entry was determined by quantifying the number of viral genomes within the cell lysate using qRT-PCR. B) rVSV/G and rVSV/EBOV-GP entry was assessed via flow cytometry. Cells were infected with rVSV/G (MOI= 0.5) for 16 hours or rVSV/EBOV-GP (MOI= 20) for 12 hours, after which cells were harvested, fixed, and analyzed for GFP fluorescence. To observe cell-cell viral spread, cells were infected with rVSV/G (MOI= 0.005) (C) or rVSV/EBOV-GP (MOI= 0.01) (D). Viral spread was monitored by GFP fluorescence over time, and 3 representative images were taken at the indicated time points (magnification, ×20). E) To assess viral replication cells were infected with rVSV/G (MOI 0.01) or rVSV/EBOV-GP (MOI 0.05) and viral genome copies were quantified in the cell lysates 12 hours (rVSV/G) or 24 hours (rVSV/EBOV-GP) post infection. Values shown are averages of at least three independent experiments ± SEM.

In order to better visualize rVSV/EBOV-GP cell–cell spread in HAP1 and ΔXKR8 cells, we plated and infected cells similarly to our multistep growth curves in Fig. 2A and B, and monitored the cellular production of virus-encoded GFP via fluorescence microscopy (Fig. 3C and D). The earliest time points after infection revealed comparable numbers of GFP-producing (GFP+) cells, confirming that the initial entry of rVSV/EBOV-GP viral stock into ΔXKR8 cells was uninhibited. However, we observed fewer GFP+ rVSV/EBOV-GP-infected ΔXKR8 cells over time (particularly at 36 and 48 hours following infection), indicating a cell-cell spread defect. Late in infection, the level of GFP+ ΔXKR8 cells reached those of HAP1 cells, mirroring the virus spread demonstrated in the growth curves (Fig. 2A). As expected, rVSV/G propagated through HAP1 and ΔXKR8 GFP with similar efficiencies as was demonstrated by no detectable differences in the level of GFP+ cells in either cell type.

We next assessed the ability of ΔXKR8 cells to support viral replication. While initial entry of our rVSV particles is executed by PS and/or the viral glycoprotein, the remaining steps in the viral replication cycle are mediated solely by VSV machinery. Because rVSV/G multi-step replication was only slightly impacted by XKR8 deletion (Fig 2B), we did not expect differences in post-entry replication stages. We quantified genome copy numbers in cell lysates 12◻hours following rVSV/G or rVSV/EBOV-GP infection and observed no significant differences in the number of genomes produced by either cell line (Fig. 3E).

### ΔXKR8 cells decrease rVSV budding efficiency

The final step in rVSV replication is virion budding at the plasma membrane, resulting in cell-free infectious particles. We produced rVSV labeled with nano-luciferase to easily quantify the amount of particles released into cell supernatants versus those inside each cell line by measuring nano-luciferase activity. Surprisingly, ΔXKR8 cells released significantly fewer rVSV-MnLuc/G and rVSV-MnLuc/EBOV-GP particles into the supernatant (Fig 4A). Using immunoblot analysis, we observed similar levels of rVSV matrix (M) protein levels in parental and ΔXKR8 cell lysates after one round of replication; however, the levels of VSV-M present in the supernatant further suggested that rVSV particle budding in ΔXKR8 cells is inefficient (Fig. 4B). Despite evidence of defective viral budding in the absence of XKR8, this defect only slightly altered rVSV/G replication in ΔXKR8 cells (Fig. 2B); thus, it does not completely account for the dramatic EBOV-GP-specific replication delay (Fig 2A).

**Figure 4.**
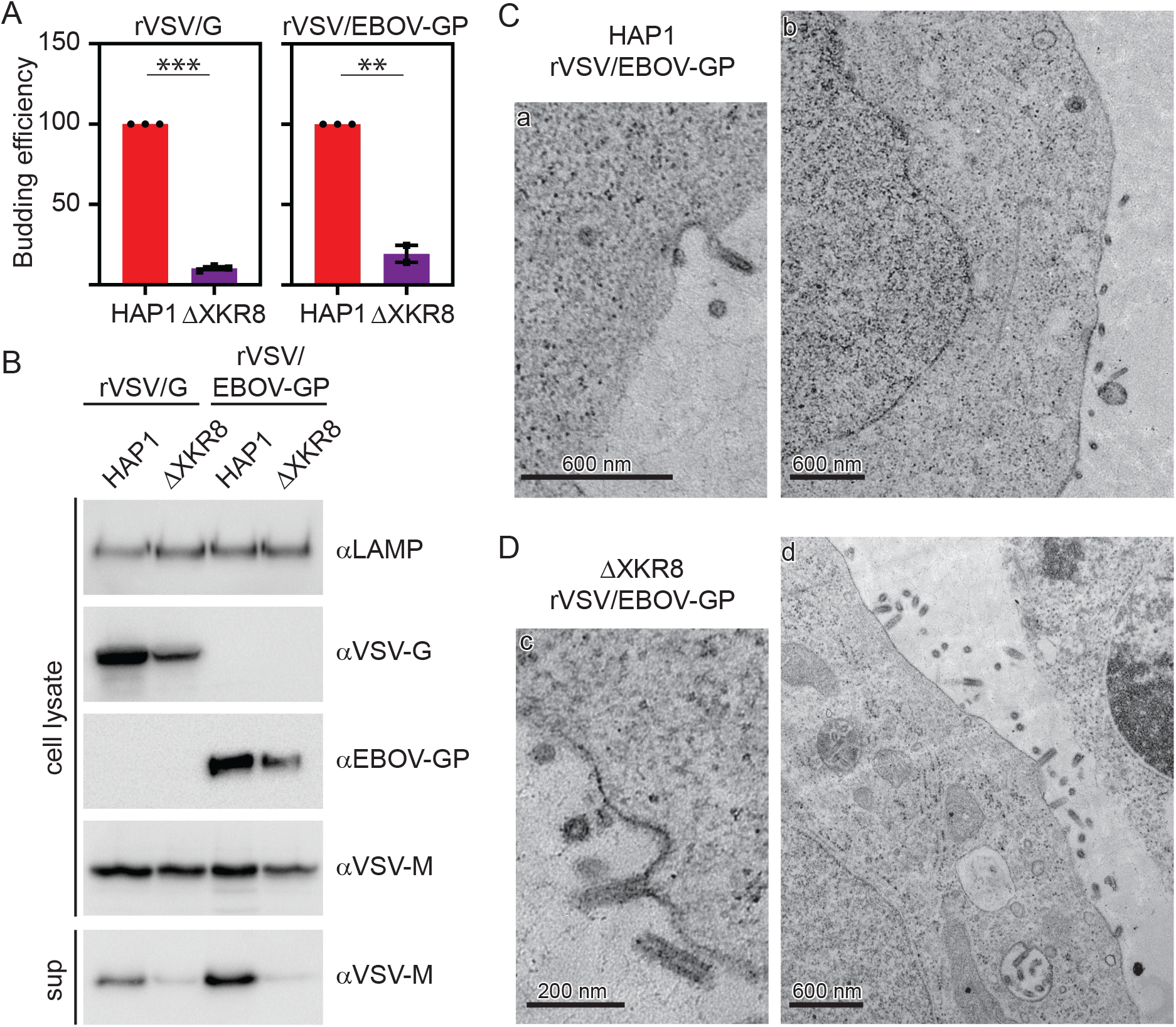
XKR8 is required for efficient rVSV budding. A) Cells were infected with rVSV-MnLuc/G (MOI= 0.1) or rVSV-MnLuc/EBOV-GP (MOI= 0.25) for 12 hours, after which cell lysate and supernatant nano-luciferase (nLuc) activity was measured. Budding efficiency was calculated by dividing supernatant nLuc activity by lysate nLuc activity, and values were normalized to HAP1 budding efficiency. B) Immunoblot probing for VSV M and VSV G or EBOV GP in rVSV/G- or rVSV/EBOV-GP-infected HAP1 and ΔXKR8 cell lysates and supernatants after one round of replication. LAMP-1 levels are shown as a cell lysate loading control. Transmission electron microscopy analysis of rVSV/EBOV-GP particles budding from the surfaces of HAP1 cells (C) or ΔXKR8 cells (D). Images were captured at the following magnifications: a: 6000x, b: 5000x, c:15000x, and d: 6000x. Values shown are averages of at least three independent experiments ± SEM, ** p<0.01, *** p<0.001, student’s t-test.

In order to visualize budding rVSV/EBOV-GP particles, we performed transmission electron microscopy on infected HAP1 (Fig. 4C) and ΔXKR8 (Fig. 4D) cells. While it was consistently difficult to locate budding events in HAP1 cells, presumably because particles were released into cell supernatant (68), we could find many occurrences of particles budding from ΔXKR8 cells. In many instances it appeared that budded rVSV/EBOV-GP particles were immobilized on the surface of the ΔXKR8 cells. While we did not observe an obvious defect in particle-membrane fission, more detailed images are needed to confirm whether the particles remaining on the surface of ΔXKR8 cells are completely separated from the plasma membrane.

### ΔXKR8-produced rVSV/EBOV-GP virions are less infectious and contain lower levels of PS

We hypothesized that rVSV/EBOV-GP replication was delayed in ΔXKR8 cells due to low levels of virion surface PS, which would decrease particle infectivity via PS receptors. In order to address specific infectivity, we produced rVSV/G and rVSV/EBOV-GP in parental and ΔXKR8 cells and compared viral genome numbers to the number of plaque-forming units in each sample (Fig. 5A). This ratio determines the number of viral genomes required to produce one plaque; therefore, a lower specific infectivity value means particles are more infectious, whereas high values suggest that many more viral particles are required to establish infection. rVSV/G specific infectivity was highly efficient and unaffected by the absence of XKR8; for every five particles detected via qPCR, one particle was infectious. For rVSV/EBOV-GP, 1 in 15 HAP1-made rVSV/EBOV-GP particles were infectious, and even fewer were infectious when virus was produced in ΔXKR8 cells (1 in 53 ΔXKR8 particles were infectious). rVSV/EBOV-GP particles were generally not as infectious as rVSV/G particles, which may be attributed to less efficient incorporation of non-native EBOV-GP into VSV particles.

**Figure 5.**
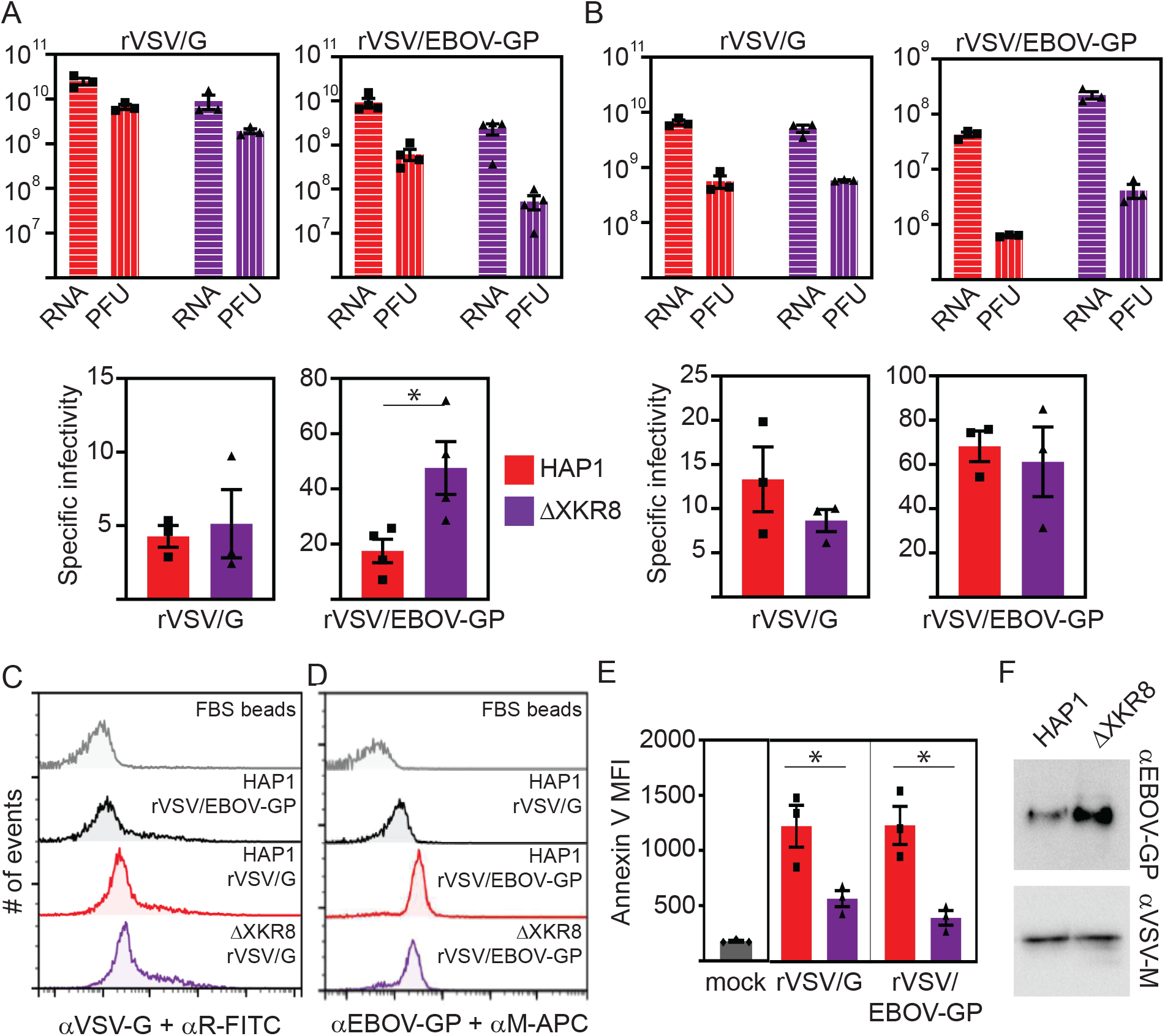
Knocking out XKR8 reduces rVSV/EBOV-GP specific infectivity and PS levels. A) rVSV/G and rVSV/EBOV-GP were propagated in HAP1 and ΔXKR8 cells for 24 hours. Supernatants were collected, filtered, and viral genomes were quantified and titrated via plaque assay (top panels). Viral specific infectivity values were determined by calculating the ratio of viral genome copy numbers to infectious particle numbers (PFU/mL) for each sample (bottom panels). B) HAP1-made or ΔXKR8-made rVSV/G or rVSV/EBOV-GP collected from part (A) were passaged once in Vero cells (rVSV/G MOI= 0.01; rVSV/EBOV-GP MOI= 0.05). After 24 hours, the supernatants were collected, and specific infectivity values were quantified. Equal amounts of rVSV/G and rVSV/EBOV-GP particles produced in HAP1 and ΔXKR8 cells were attached to aldehyde-sulfate latex beads and immunostained for VSV G (C) or EBOV GP (D). E) Viral particles were stained for surface PS levels with AnV and MFIs were quantified. F) Equivalent levels of rVSV/EBOV-GP particles produced by HAP1 and ΔXKR8 cells were analyzed for VSV M and EBOV GP protein content by immunoblot. Values shown are averages of at least three independent experiments ± SEM, *p<0.05, student’s t-test.

In order to demonstrate that reduced rVSV/EBOV-GP infectivity was the direct result of ΔXKR8-made viral envelopes, we passaged the samples generated in Fig. 5A through Vero cells and determined the specific infectivity of the viral progeny (Fig. 5B). The rVSV/G and rVSV/EBOV-GP progeny specific infectivity values were similar with respect to viral GP after Vero passage, regardless of the original producing cell lines. These data support our hypothesis that host-cell scramblase deletion reduces rVSV/EBOV-GP infectivity due to alterations in the viral envelope and that these alterations are irrelevant for rVSV/G, a virus that does not utilize apoptotic mimicry.

In order to confirm that viral particles produced in ΔXKR8 cells contain lower levels of outer leaflet PS, we coated latex beads with rVSV particles produced in parental and ΔXKR8 cells and examined viral envelope AnV-PacBlue fluorescence intensity using flow cytometry. Incubating beads with 1.5 × 10^9^ genomes of rVSV/G or rVSV/EBOV-GP ensured that beads were uniformly and completely coated in viral particles, indicated by robust G or GP staining (Figs. 5C and D). Producing rVSV in ΔXKR8 cells significantly reduced outer leaflet PS levels in a GP-independent manner, resulting in an approximately two-fold decrease in the AnV mean fluorescence intensity on rVSV/G particles and a three-fold decrease on rVSV/EBOV-GP particles (Fig. 5E).

We further analyzed parental and ΔXKR8-made rVSV/EBOV-GP for GP incorporation, because a defect in envelope GP levels could also reduce particle specific infectivity during viral fusion and genome release. Supernatant VSV genome copy numbers were quantified using qRT-PCR, and normalized volumes were subjected to immunoblot analysis for VSV M and EBOV GP (Fig. 5F). While we previously observed lower glycoprotein levels within the cells (Fig. 4B), we consistently found higher GP:matrix levels in ΔXKR8-produced rVSV/EBOV-GP particles. This data suggests that the reduced specific infectivity observed in ΔXKR8-produced rVSV/EBOV-GP particles was not due to poor GP incorporation.

### EBOV VLPs require XKR8 for externalized envelope PS

While the rVSV system enables us to monitor EBOV-GP-mediated entry in a replication-competent viral vector, the rVSV particle is morphologically distinct from authentic EBOV. rVSV is also a strong inducer of apoptosis and may activate XKR8 more profoundly compared to EBOV. Therefore, we employed EBOV virus-like particles (VLPs) to further examine the role of XKR8 in EBOV replication. The Ebola matrix protein VP40 drives VLP production, therefore to quantify VLP production and budding, we fused VP40 to either nano-luciferase or GFP to produce nLuc-VLPs and GFP-VLPs. HAP1 versus ΔXKR8 VLP budding efficiencies were measured using nLuc-VLPs. VLP budding efficiency was significantly reduced in ΔXKR8 cells compared to HAP1 cells (Fig 6A). Because XKR8 is activated by caspase cleavage, we also examined VLP budding in HAP1 cells treated with pan-caspase inhibitor Z-VAD-FMK. Preventing XKR8 activation with Z-VAD-FMK reduced VLP budding almost as effectively as a full XKR8 gene knockout (Fig. 6B). Next, we over expressed XKR8 by transfecting HAP1 and ΔXKR8 cells with an XKR8 FLAG-tagged construct. nLuc-VLP budding efficiency increased 4-fold with the addition of XKR8_FLAG_ in both HAP1 and ΔXKR8 cells, and the increase was reversed by Z-VAD-FMK, suggesting XKR8 activation via caspase cleavage inhibits nLuc-VLP budding (Fig. 6C). We also over-expressed XKR8 in Vero cells and observed a profound increase in budding efficiency, indicating that XKR8 activity is required for budding in multiple cell types (Fig. 6D). Nanbo et al. generated EBOV VLPs in 293T cells producing a shRNA targeting XKR8 and found that without XKR8, EBOV VLPs contained less surface PS and were internalized poorly into Vero E6 cells (48). To confirm that this VLP effect was not 293T-specific, we made GFP-VLPs in our HAP1 and knocked-out ΔXKR8 cells with or without exogenous XKR8_FLAG_ and evaluated VLP EBOV GP and PS surface levels using flow cytometry (Fig. 6E and F). GFP-VLP GP incorporation was not affected by the absence of XKR8, however, over-expressing XKR8 reduced VLP GP levels (Fig. 6E). Meanwhile, GFP-VLPs made in ΔXKR8 cells displayed significantly lower PS levels in comparison to HAP1 cells (Fig. 6F). When we over-expressed XKR8, we observed increased GFP-VLP surface PS levels, confirming the importance of XKR8 activity in EBOV surface PS.

**Figure 6.**
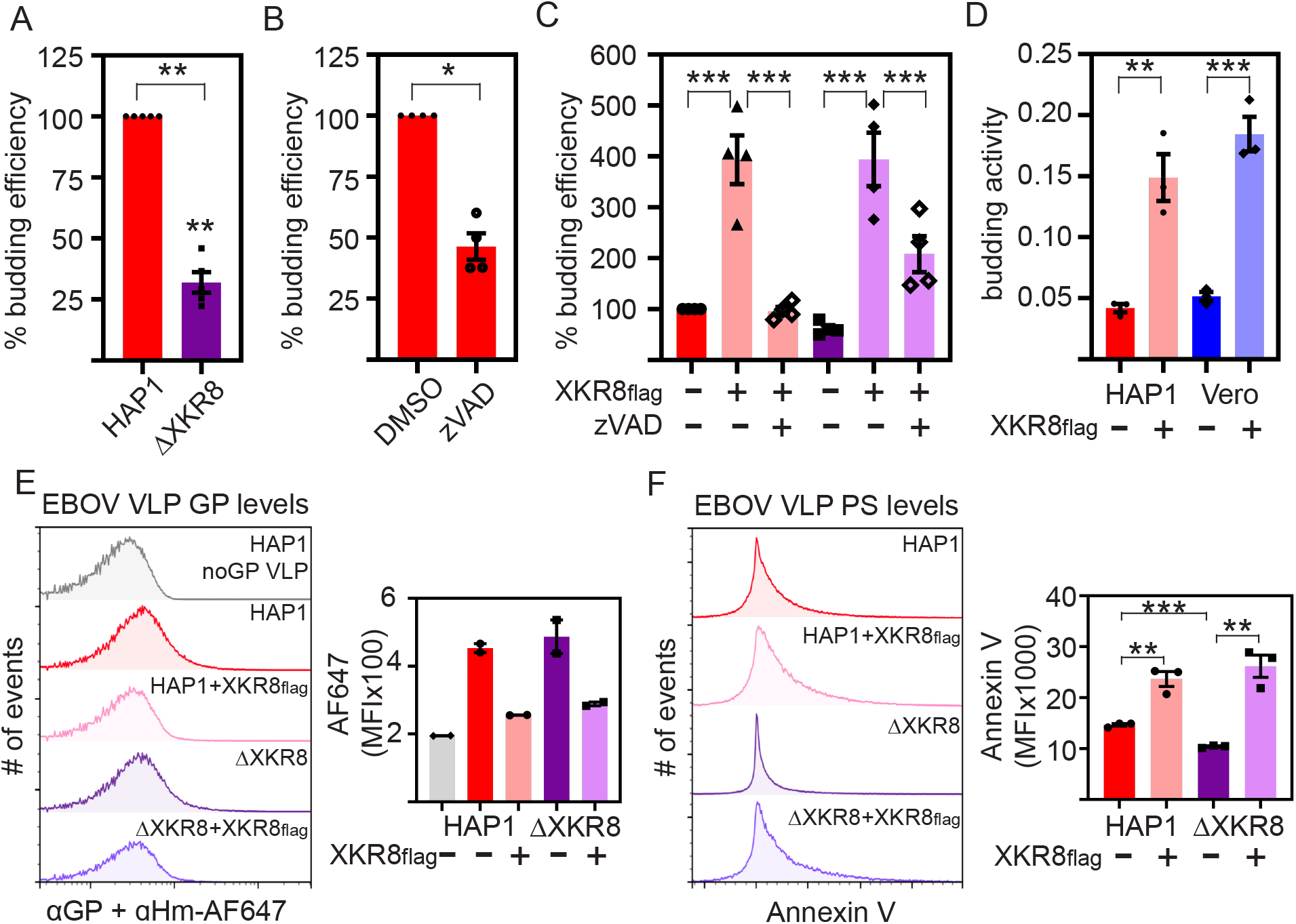
XKR8 activity is required for efficient EBOV VLP budding and particle PS levels. A) HAP1 and ΔXKR8 cells were transfected with EBOV VLP components NP, GP, and VP40-nano-luciferase and budding activity was measured 24 hours following transfection; values were normalized to HAP1 budding efficiency. B) nLuc-VLPs were produced in HAP1 cells in the presence of DMSO or pan-caspase inhibitor Z-VAD-FMK (20μM). C) nLuc-VLPs were produced in HAP1 and ΔXKR8 cells with or without exogenous XKR8_FLAG_, and in the presence of either DMSO or Z-VAD-FMK. D) nLuc-VLPs were produced in HAP1 and Vero cells with or without exogenous XKR8_FLAG_. nLuc-VLP budding efficiencies were quantified. E) EBOV GP levels on VLPs produced in HAP1 and ΔXKR8 cells with or without exogenous XKR8_FLAG_, VLPs were produced without GP for a negative control. Anti-EBOV-GP/anti-human AF647 fluorescence values for GFP+ VLPs are represented as histograms (left panel) and average MFIs (right panel). F) GFP-VLPs were also stained for surface PS using AnV. AnV-PE fluorescence values of GFP+ VLPs are represented as histograms (left panel) and average MFIs (right panel). Values shown are averages of at least three independent experiments ± SEM, * p<0.05, ** p<0.01, *** p<0.001, student’s t-test.

## Discussion

Although previous paradigms suggest that all viral GPs orchestrate enveloped viral attachment to host cells in a lock-and-key-like manner, evidence now suggests that less-specific mechanisms of virion attachment can be utilized to facilitate internalization. Attachment factors such as lectins and PS receptors effectively bind and internalize viral particles without specific GP interaction (12, 20, 21, 23, 24). In order to interact with cellular PS receptors, viral particles must contain PS in the outer leaflet of their envelopes. However, normal-state cells strictly sequester PS in the inner leaflet of the plasma membrane. Therefore, it has been speculated in previous studies on apoptotic mimicry that viruses budding from the plasma membrane trigger cellular PS externalization through endogenous pathways, such as apoptosis, to obtain surface PS-rich viral envelopes. To determine whether cellular scramblases, proteins required to move phospholipids between the inner and outer leaflets of cell membranes, are involved in viral PS acquisition, we monitored EBOV-GP-mediated spread in CRISPR knockout HAP1 cells deficient in XKR8 or TMEM16F scramblases. Here, we demonstrate that the activity of XKR8, a transmembrane scramblase required for apoptotic cell PS externalization, is also required for elevated PS levels on rVSV/EBOV-GP and EBOV VLP envelopes. The EBOV envelope PS levels achieved by XKR8 activation are also critical for rVSV/EBOV-GP infectivity.

Both our group and Nanbo et al. primarily attribute increased EBOV envelope surface PS to XKR8 activation, which requires cleavage by apoptosis effector caspase-3. EBOV-induced apoptosis is considered to be controversial, owing to a 2013 study showing weak caspase activation and variable PS exposure in EBOV-infected Vero E6 cells (69). Nanbo et al. discovered similarly limited XKR8 activation in VLP-transfected Vero E6 cell lysates and observed only minor plasma membrane PS scrambling when VP40 and GP were co-expressed; however, they reported significant XKR8 activation in EBOV VLP envelopes (48). Weak cellular EBOV-induced XKR8 activation may be cell-type-specific, as Nanbo et al. also found XKR8 protein uncharacteristically confined to intracellular vesicles in Vero E6 cells unless cells simultaneously produced EBOV GP. Studies on alternative cell types HEK293 and Huh7 show robust EBOV virus or EBOV protein-induced PS externalization in plasma membranes (34, 49). We demonstrate that drug-induced XKR8-mediated apoptosis causes significantly higher HAP1 surface PS levels, indicating that XKR8 is sufficiently trafficked to the plasma membrane in these cells without viral proteins.

Younan et al. examined WT EBOV PS acquisition in Huh7 liver cells producing shRNAs targeting TMEM16F or XKR8 (49). This group observed widespread elevated PS on EBOV-infected cells or GP/VP40-transfected cells, which was partially reduced on Huh/shXKR8 cells and significantly reduced on Huh/shTMEM16F cells. They treated EBOV-infected Huh7 cells with either a calcium chelator or a calcium-dependent scrambling inhibitor for 48◻h and reported fewer virus-infected cells and significantly less PS scrambling on cell surfaces. Therefore, they named TMEM16F the primary mediator of EBOV surface PS during peak infection. Their model describes a process by which EBOV-GP induces ER stress in Huh7 cells, causing the release of ER calcium stores into the cell cytoplasm, resulting in calcium-dependent TMEM16F activation, plasma membrane PS scrambling, and WT EBOV particles with increased outer leaflet PS. They suggested that XKR8 may marginally contribute to cellular and viral enhanced PS levels during late-stage cellular apoptosis. We found that calcium-induced PS scrambling is TMEM16F- and XKR8-independent in HAP1 cells, as we observed high levels of annexin staining in cells completely lacking TMEM16F or XKR8 after treatment with calcium ionophore A23187. HAP1 cells produce TMEM16D, another calcium-dependent scramblase from the TMEM16 family, which may facilitate PS scrambling either exclusively or in conjunction with TMEM16F (70). Predictably, eliminating TMEM16F activity did not alter rVSV/EBOV-GP viral spread in HAP1 cells. Therefore, these data do not support a role for TMEM16F in this system; however, calcium-activated scramblase activity may be important in specific cell types.

While the two aforementioned studies identified scramblases required to facilitate EBOV surface PS using shRNA knockdown cells, neither study reported budding defects. The Nanbo et al group performed extensive particle purification and concentration (48), which may have prevented head-to-head comparison of the produced VLP yields. Younan et al infected Huh7/shXKR8 and Huh7/sh control cells with EBOV-GFP, monitored cell GFP production over time, and found no discrepancies in EBOV-GFP spread (49). However, immunoblot analysis of their Huh7/shXKR8 cell line showed a modest decrease in XKR8 production in comparison to wild-type Huh7 cells, which may have been insufficient to affect EBOV replication.

We postulate that EBOV induces XKR8 activation and widespread cell apoptosis in a highly coordinated manner with viral particle assembly and budding, which maximizes both viral budding efficiency and virion infectivity. Studies by Adu-Gyamfi et al. showed that EBOV VP40 specifically binds to inner leaflet plasma membrane PS, and this interaction is required for efficient VP40 trafficking, oligomerization, and egress (34). In the same study, they also reported VP40-induced PS scrambling in transfected HEK293 cells. While seemingly counterintuitive, Adu-Gyamfi et al. suggested that VP40 dimers oligomerizing at the plasma membrane inner leaflet may eventually stabilize and release PS, and that subsequent enzyme-mediated PS externalization would promote particle egress by inducing positive membrane curvature. The EBOV VLP budding defects that we observed in ΔXKR8 cells support this proposed mechanism. Particles assembling at the inner leaflet of ΔXKR8 cells may be unable to bud efficiently because PS externalization is inhibited. In our rVSV system, particle assembly and budding are mediated by VSV proteins, yet we still observed significantly reduced budding in ΔXKR8 cells, suggesting that both VSV and EBOV particle budding are enhanced with XKR8 activity.

Evaluating the importance of XKR8 in more biologically relevant systems will be critical for characterizing the EBOV apoptotic mimicry mechanism. Producing BSL-4 EBOV in complete scramblase knockout cells will enable us to fully define XKR8’s contribution to EBOV envelope surface PS and infectivity. Research investigating EBOV-induced effects on specific cellular apoptosis and calcium effectors upstream of scramblases may provide additional clarity on the driving force behind EBOV envelope PS and the potential interplay between VP40 and PS scramblases. Furthermore, future experiments more closely examining VSV and EBOV budding in ΔXKR8 cells will provide important insights into viral budding mechanics.

## Acknowledgments

We that the CVM Cytometry Core Facility, especially James Barber for technical assistance, members of the Brindley lab for helpful comments on the manuscript.

This work was supported by the National Institutes of Health (R01 AI139238).

